# Exogenous ephrin-A3 administration restores vaginal epithelial barrier function in progestin-treated mice

**DOI:** 10.1101/2024.10.29.620915

**Authors:** Mohan Liu, Rodolfo D Vicetti Miguel, Kristen Aceves, Thomas L Cherpes

**Author notes:** Corresponding Author: Rodolfo Vicetti Miguel, MD, 5111 Pelotonia Research Center, 2255 Kenny Rd, Columbus, OH USA 43210.

## Abstract

Desmosomes are junctional complexes that confer mechanical strength and enhance epithelial barrier function at mucosal surfaces by anchoring intermediate filaments to plasma membrane. While these roles are less explored in vaginal vs. cutaneous epithelium, we previously reported that treating mice with the progestin depot medroxyprogesterone acetate (DMPA) reduces vaginal epithelial levels of the desmosomal cadherins desmoglein-1 (DSG1) and desmocollin-1 (DSC1) and weakens vaginal epithelial barrier function. We also showed these effects were avoided by treating mice with DMPA and a conjugated equine estrogen vaginal cream. The current investigation further explored the effects of sex steroids on vaginal epithelial integrity, identifying ephrin-A3 (EFNA3) as a key regulator of desmosomal cadherin gene expression. We observed topical administration of recombinant EFNA3 (rEFNA3) promotes vaginal DSG1 expression in a biphasic dose-dependent manner and partially reverses the loss of vaginal epithelial barrier function induced by DMPA treatment. Consistent with this effect, morbidity and mortality elicited by genital herpes simplex virus type 2 infection were delayed, but not prevented, in mice administered DMPA and rEFNA3 vs. DMPA and vehicle. Together, these studies identify EFNA3 as an important regulator of desmosomal function in vaginal epithelium and improve current understanding of sex steroid-mediated mechanisms that control vaginal epithelial barrier function.

## INTRODUCTION

Mucosal surface epithelial cells are connected by tight junctions, adherens junctions, and desmosomes that restrict paracellular migration of microbes and other luminal content ^1–3^. While all three junctional complexes promote epithelial cell-cell adhesion, desmosomes promote barrier function and confer mechanical strength by anchoring intermediate filaments to plasma membrane ^4^. As desmosomes resist mechanical stress with strong epithelial attachments, they are also called anchoring junctions. These junctions are composed of the transmembrane cadherin proteins desmoglein and desmocollin, an associated cytoskeletal adaptor complex formed by plakoglobin and plakophilin, and desmoplakin ^5^. The importance of desmosomes for conserving epithelial health is evidenced by the profound loss of cutaneous epithelial barrier function and enhanced susceptibility to recurrent infection in individuals genetically deficient in desmoglein-1 (DSG1) ^6–8^. Likewise, autoantibodies against DSG1 cause the severe blistering characteristic of the potentially fatal skin disease pemphigus ^9^.

Supporting a role for desmosomes at other sites of stratified squamous epithelium, cervicovaginal mucosa is often affected in individuals with pemphigus vulgaris ^10^. Clinical studies also identified significantly lower levels of DSG1 and desmocollin-1 (DSC1) in ectocervical epithelium from women that initiated use of the synthetic progestin depot medroxyprogesterone acetate (DMPA) for contraception ^11, 12^. Indicating this loss of epithelial integrity weakens barrier protection, we also visualized deeper penetration of the low-molecular-mass molecule (LMM) lucifer yellow (LY) into ectocervical epithelium of women that initiated use of DMPA ^11^. Compared to mice in the estrus stage of the estrous cycle, DMPA-treated mice likewise displayed loss of genital DSG1 and DSC1 levels, impaired genital epithelial barrier function, and enhanced susceptibility to genital infection with herpes simplex virus type 2 (HSV-2) ^11, 13^. Because DMPA disrupts the hypothalamic-pituitary-ovarian (HPO) axis and produces a hypoestrogenic state by inhibiting ovarian follicle maturation and 17 β-estradiol synthesis, our results suggested that estrogen (E) promotes genital epithelial integrity and barrier function. Congruent with this effect of endogenous E, DMPA-treated mice were made fully resistant to genital HSV-2 infection by intravaginal (ivag) administration of conjugated equine estrogen cream (CEE) (i.e., Premarin®) ^11^. We analogously observed that exogenous E reverses DMPA-mediated loss of genital epithelial integrity and barrier function in nonhuman primates ^14, 15^.

Indication that exogenous and endogenous sex steroids regulate genital epithelial health provided rationale in the current investigation to further explore links between HPO axis disruption and loss of genital epithelial integrity and barrier function. Our initial study used genome-wide expression profiling of vaginal tissue from mice in estrus (the estrous cycle stage dominated by E effects) and DMPA-treated mice to define regulators of desmosomal cadherin gene expression. After identifying the ephrin-A3 (EFNA3) pathway as an important regulator of sex steroid-mediated effects in genital epithelium, we used additional studies to define the role of this pathway in promoting genital epithelial integrity and barrier function.

## MATERIALS and METHODS

### Mice and treatments

Study procedures were approved by Institutional Animal Care and Use Committees at Stanford University and Ohio State University prior to acquiring 8–10-week-old female C57BL/6 mice from Jackson Laboratory (Bar Harbor ME). Using defined methods, estrous cycle stage was diagnosed by light microscope exam of crystal violet-stained cells from vaginal lavage ^16^. As indicated, mice were subcutaneously (SQ) injected with 30 mg/kg of DMPA (Mylan Institutional, Rockford IL). Commercially available recombinant mouse EFNA3 (rEFNA3) was purchased as Fc fusion (R&D Systems, Minneapolis MN) or His-tagged proteins (SinoBiological, Wayne PA). Where indicated, recombinant mouse IgG2A Fc protein provided a Fc control (R&D Systems). His-tagged versions of rEFNA1, 2, 4 and 5 (SinoBiological) were also obtained and rEFNA proteins administered to mice sedated with isoflurane inhalation anesthetic (Dechra Veterinary Products, Overland Park KS). Prior to treatment, vaginal fluid was absorbed by atraumatic insertion of cellulose eye spears (Beaver Medical, Waltham VA). Sedated mice were intravaginally administered indicated doses of rEFNA3 (SinoBiological, Wayne PA) reconstituted in sterile phosphate buffered saline (PBS) (Mediatech Inc., Manassas VA). Mice were placed in the dorsal recumbent position for 3 hours and recovered from anesthesia. In experiments that investigated EFNA promiscuity, rEFNA1, 2, 3, 4, or 5 (all His-tagged) were prepared and administered using methods identical to those detailed above.

### Genome-wide gene expression profiling

For gene expression profiling, vaginal tissue from estrus-stage (n=3) and DMPA-treated (n=3) mice were placed in RNAlater RNA Stabilization Reagent and total RNA extracted using the RNeasy Lipid Tissue Kit (both Qiagen, Hilden Germany). RNA concentrations were measured via NanoDrop spectrophotometer (Thermo Fisher Scientific, Pittsburgh PA) and RNA Integrity Index (RIN) values defined via Agilent Bioanalyzer (Agilent Technologies, Santa Clara CA) (RIN values were routinely > 8). Genome-wide gene expression profiling was performed using the Affymetrix GeneChip® Mouse Transcriptome Assay 1.0 and CEL files imported to the Transcriptome Analysis Console (Applied Biosystems, Santa Clara CA). After SST-RMA normalization, differentially expressed genes in tissue from estrus-stage vs. DMPA-treated mice were identified using limma (Bioconductor). The significant limit for functional enrichment analysis was defined by a false discovery rate-adjusted *P* value < 0.05 and a fold change in gene expression of at least <-2 or >2. Differentially expressed genes were analyzed using QIAGEN’s Ingenuity Pathway Analysis (IPA) software (QIAGEN, Redwood City CA). The core analysis function of IPA was used to interpret the data in the context of biological processes, pathways, and networks. Genes that met the indicated threshold of fold change and *P* value were uploaded into IPA along with their corresponding expression values. The core analysis identified the canonical pathways, upstream regulators, and biological functions most significantly associated with the dataset. Significance of the association between dataset and canonical pathway was measured by Fisher’s exact test. *P* values < 0.05 identified statistically significant, non-random associations. Activation z-scores were used to infer likely activation states of upstream regulators based on comparison with a model that assigns random regulation directions and absolute values of z-score >2 were considered significant.

### Histology, immunohistochemistry, and immunofluorescent staining

Vaginal tissue collected 24 hours after rEFNA3 or other rEFNA treatment was fixed in 4% formaldehyde overnight before transfer to 70% reagent alcohol. Samples were paraffin-embedded, sliced into 5 μm sections, and stained with hematoxylin and eosin (H&E). For DSG1 immunohistochemistry (IHC), 4 μm paraffin-embedded sections were placed on Superfrost Plus slides to dry overnight and stepwise deparaffinization performed with 5-minute incubations in xylene and decreasing ethanol concentrations. For antigen retrieval, sections were placed 20 minutes into dishes containing Target Retrieval Solution (Dako, Santa Clara CA) preheated to 90°C, cooled, rinsed with 1X Tris-buffered saline (TBS), and incubated for 15 minutes in PBS containing 3% (v/v) hydrogen peroxidase to block endogenous peroxidase activity. A hydrophobic pen mark was made to denote tissue location and slides placed in TBS containing 0.1% (v/v) Tween-20. Slides were air dried and blocked for 20 minutes with normal horse serum (Vector Laboratories, Newark, CA) and anti-DSG1 primary antibody (ab124798) (Abcam, Waltham, MA) (0.125 µg/mL) in Dako antibody diluent for 30 minutes. Slides were washed in TBS with 0.1% (v/v) Tween-20 and incubated for 30 minutes with donkey anti-rabbit secondary antibody (Vector, Newark, CA). DAB substrate (Dako) was added for 3 minutes and slides rinsed in distilled water. Slides were incubated 30 seconds in Harris Hematoxylin (Leica Biosystems, Deer Park IL) and rinsed 30 seconds in distilled water containing 0.1% NH_4_OH (v/v). Tissue was dehydrated with increasing ethanol and xylene concentrations and whole slide imaging performed at 20x magnification. Immunofluorescence (IF) staining was performed as previously described ^16^ using primary antibodies anti-DSG1 (Clone EPR6766(B), Abcam) and anti-EFNA3 (rabbit polyclonal, SinoBiological) antibodies. Slides were examined by Nikon A1R confocal microscope, and relative intensity of staining determined using Fiji, a distribution of ImageJ software (NIH).

### RNA isolation and quantitative real-time PCR (qRT-PCR)

For qRT-PCR, vaginal tissue was immediately immersed in RNAlater and RNA isolated with RNeasy Lipid Tissue Kits. RNA concentrations were measured via NanoDrop^TM^ One/One^C^ Microvolume UV-Vis Spectrophotometer (Thermo Fisher Scientific, Rockford, IL), and 500 ng of RNA used to generate cDNA using SuperScriptTM IV VILOTM master mix and ezDNaseTM enzyme (Thermo Fisher Scientific, Rockford IL). *Efna3* (Mm07296616_m1) and *Dsg1a* (Mm00809994_s1) relative gene expression were evaluated using pyruvate carboxylase (Mm00500992_m1) as housekeeping gene (Life Technologies, Carlsbad, CA), and qRT-PCR performed using a QuantStudio 3 (Applied Biosystems, Thermo Fisher Scientific, Rockford, IL). Results were analyzed using the QuantStudio^TM^ Design & Analysis Software (Thermo Fisher Scientific, Rockford, IL).

### Genital epithelial permeability

Using a previously described permeability assay, epithelial barrier function was compared in excised vaginal tissue from mice receiving DMPA vs. DMPA and rEFNA3 (or rEFNA2) 24 hours after rEFNA administration ^11, 16^. In brief, mice were sedated with 1.8 mg of ketamine hydrochloride (JHP Pharmaceuticals, Rochester MI) and 0.18 mg of xylazine (Lloyd Laboratories, Shenandoah IA) and vaginal fluid absorbed with cellulose eye spears prior to atraumatic intravaginal administration of 10 μL of PBS containing 10 μg of Lucifer Yellow CH, lithium salt (LY) (MW=457.2 Da) and 12.5 μg of Texas Red-dextran (DR) (MW=70,000) (both Thermo Fisher Scientific). Mice were placed in dorsal recumbency for 45 minutes before euthanasia and vaginal tissue excised, fixed overnight in 4% buffered formaldehyde, and vertically embedded in 6% agarose. The next day, 250 μm sections were transversely sliced using a PELCO EasiSlicer^TM^ (Ted Pella Inc., Redding, CA). Tissue blocks were counterstained with DAPI (4’, 6-diamidino-2-phenylindole, Dihydrochloride) (Invitrogen, Waltham, MA) and cover slips placed using Vectashield Mounting Media Hardset (Vector Laboratories, Newark CA). Slides were examined by Nikon A1R confocal microscope and LY molecule infiltration into vaginal epithelium (displayed in green channel images) quantified using Fiji (with values normalized to those seen in mice treated with DMPA alone).

### Genital HSV-2 infection

Mice sedated with xylazine and ketamine hydrochloride were intravaginally inoculated with 10^4^ plaque-forming units (pfu) of wild-type HSV-2 strain 333 in 10 μL of RPMI ^11, 17^. Genital pathology was monitored daily from 2-20 days post-infection (dpi) using a previously described 5-point scale (0, no infection; 1, vulvar erythema; 2, vulvar area swelling and erythema; 3, severe swelling and erythema of vulva and surrounding tissue; 4, perineal fur loss and ulceration; 5, severe genital ulceration extending to surrounding tissue). Mice were euthanized when scores were ≥ 3 or when signs of encephalopathy were observed.

### Statistical considerations

Statistical analyses were performed using Prism 10 software (GraphPad, La Jolla, CA). Ordinary one-way ANOVA with Tukey’s or Dunnett’s post hoc test were used for multiple comparisons. Kaplan-Meier survival curves and log-rank test were used to evaluate cumulative survival incidence after HSV-2 infection. *P* values ≤ 0.05 were deemed statistically significant.

## RESULTS

### DMPA decreases EFNA3 signaling in vaginal tissue

As our prior work identified that DMPA reduces genital DSG1 expression in mice, nonhuman primates, and humans ^11, 13, 14, 18^, the current study began by comparing genome-wide gene expression profiles in vaginal tissue from estrus-stage vs. DMPA-treated mice. This analysis identified significant modulation of more than 1,200 genes (Fig. 1A-C and Suppl Table S1), including genes associated with keratinocyte differentiation, skin formation and differentiation, skin barrier integrity, leukocyte migration, and immune cell adhesion (Fig.1D and Suppl Table S2). These results were consistent with our prior report of DMPA-induced loss of genital epithelial barrier integrity and barrier function and DMPA-induced genital inflammation in humans and mice ^11^. Current canonical pathway analysis also identified significant enrichment of modulated genes in pathways associated with keratinization, neutrophil degranulation, IL-10 signaling, and ephrin receptor signaling (Fig. 1E-1F and Suppl Table S3). Additional analysis identified medroxyprogesterone acetate and the proinflammatory cytokine colony stimulating factor 2 (CSF2) as upstream regulators of DMPA-mediated effects on vaginal gene expression (Fig. 1G-H and Suppl Table S4) ^19–26^.

**Figure 1.**
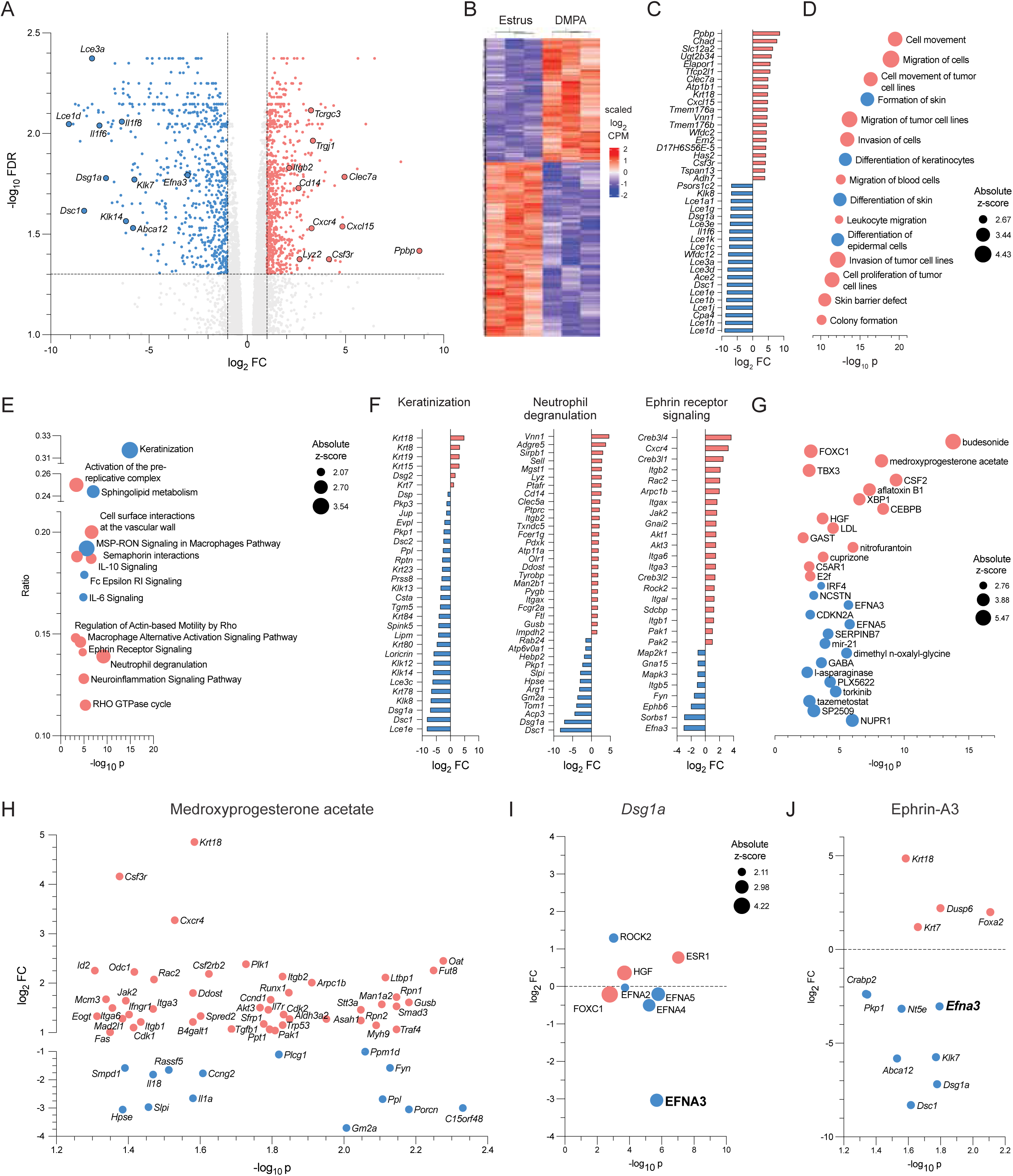
DMPA reduces DSG1 gene expression. Genome-wide gene expression profiles in vaginal tissue from estrus-stage (n=3) and DMPA-treated (n=3) mice were compared using limma and Ingenuity Pathway Analysis (IPA). A) Volcano plot identifying DMPA-modulated gene expression. B) Hierarchical clustering details differential gene expression with FDR-adjusted P value <0.05 in DMPA-treated mice and mice in estrus. C) 20 most up- and down-regulated genes in DMPA-treated mice. D) IPA identities diseases and functions associated with DMPA-modulated genes. IPA also identifies E) canonical pathways activated and inhibited by DMPA and F) genes modulated in the selected canonical pathways. G) Potential upstream regulators of DMPA effects in vaginal tissue. H) MPA-modulated gene enrichment. I) Upstream regulators of DSG1 expression. J) Genes involved in EFNA3-mediated pathways. DMPA, depot medroxyprogesterone acetate; DSG1, desmoglein 1; EFNA3, Ephrin-A3; MPA, medroxyprogesterone acetate. Upregulated genes/pathways (light red) and downregulated genes/pathways (light blue).

Congruent with our earlier reports ^11, 13, 14, 18^, genome-wide gene expression profiles that compared vaginal tissue from estrus-stage vs. DMPA-treated mice demonstrated that DMPA sharply decreases expression of the desmosomal cadherins *Dsg1a* and *Dsc1* (Fig. 1A). Exploration of possible upstream regulators targeting *Dsg1a* identified 9 potential candidates (Fig. 1I and Table 1). Among these upstream regulators, ephrin-A (EFNA) family members were well represented, with *Efna3* most sharply downregulated by DMPA treatment (Fig. 1I). Relevant to our research, EFNA family members are known to share gene expression signatures and play overlapping roles in keratinocyte differentiation, migration, and adhesion ^27–29^. In addition to *Dsg1a* and *Dsc1,* EFNA3-modulated target genes (Fig. 1J) included *Crabp2 and Abca12,* which support epidermal homeostasis and barrier function, and *Pkp1* and *Klk7,* which have direct effects on desmosomal integrity ^30,31^. Another EFNA3-modulated gene, *Nt5e*, encodes a surface enzyme known to mediate adhesion and cell signaling with fibronectin and other elements of the extracellular matrix ^32^.

**Table 1.**
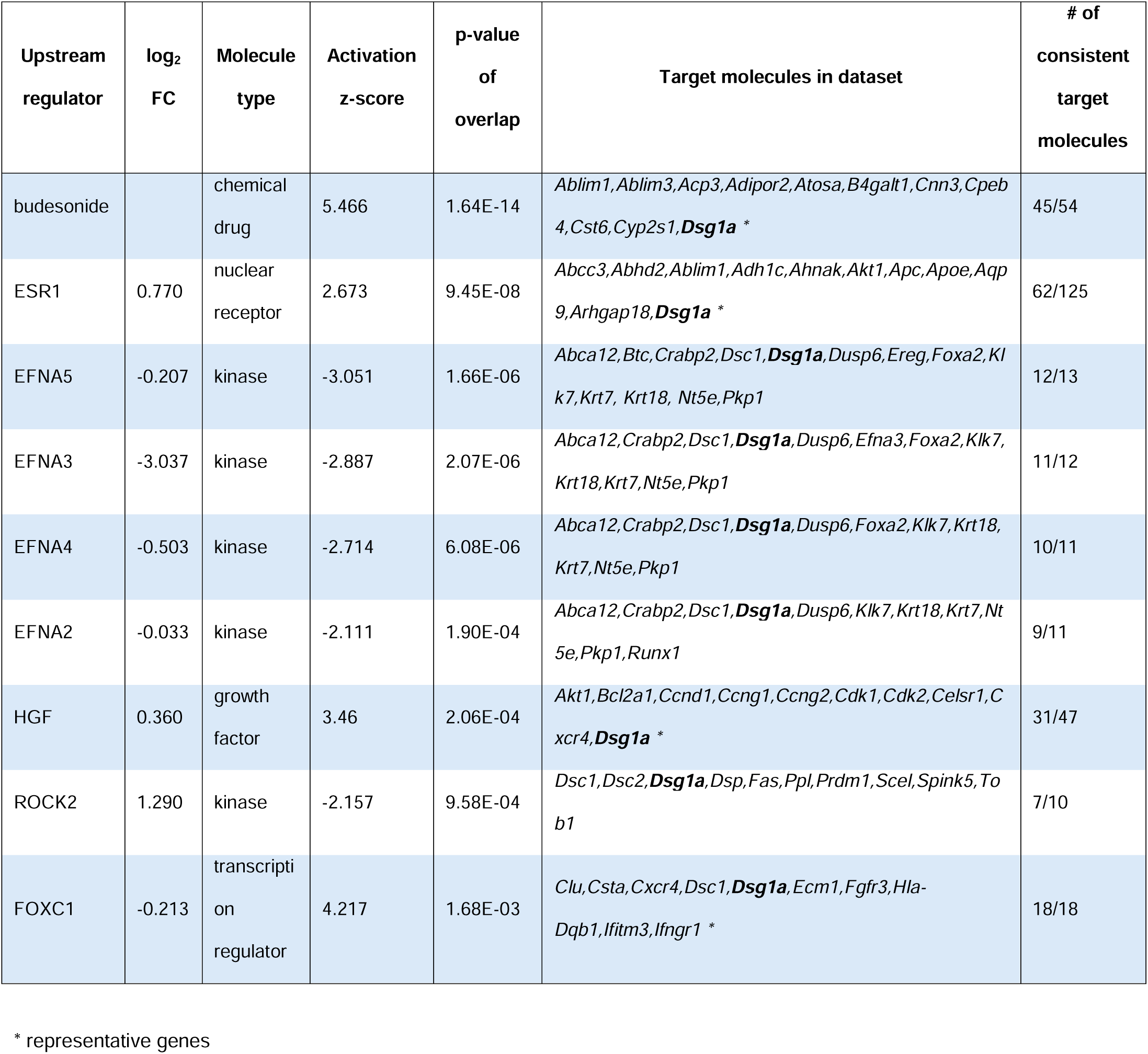
Identified upstream regulators reported to control *Dsg1a* expression.

In follow-up studies, we corroborated our microarray results with qRT-PCR and IF studies that respectively detected lower expression of the *Efna3* gene and EFNA3 protein in vaginal tissue from DMPA-treated vs. estrus-stage mice (Fig. 2A-2C). Consistent with a prior publication that showed intravaginally administered CEE (e.g., Premarin®) averts DMPA-mediated loss of genital epithelial barrier function in mice ^11^, our study identified comparable EFNA3 levels in vaginal tissue from CEE-treated and estrus-stage mice (Fig. 2A-2C). Consistent with the effects of treatment with DMPA or CEE on EFNA3 expression, we also found vaginal DSG1 levels were reduced by DMPA and restored to levels measured in estrus-stage mice by CEE (Fig. 2D, 2E). Congruent with studies that indicated EFNA-Eph signaling promotes keratinocyte differentiation in stratified squamous cutaneous epithelium ^28, 29^, current findings provide implication that EFNA3 signaling pathways in the stratified squamous epithelium of the mouse vagina increase desmosomal cadherin levels and promote epithelial integrity.

**Figure 2.**
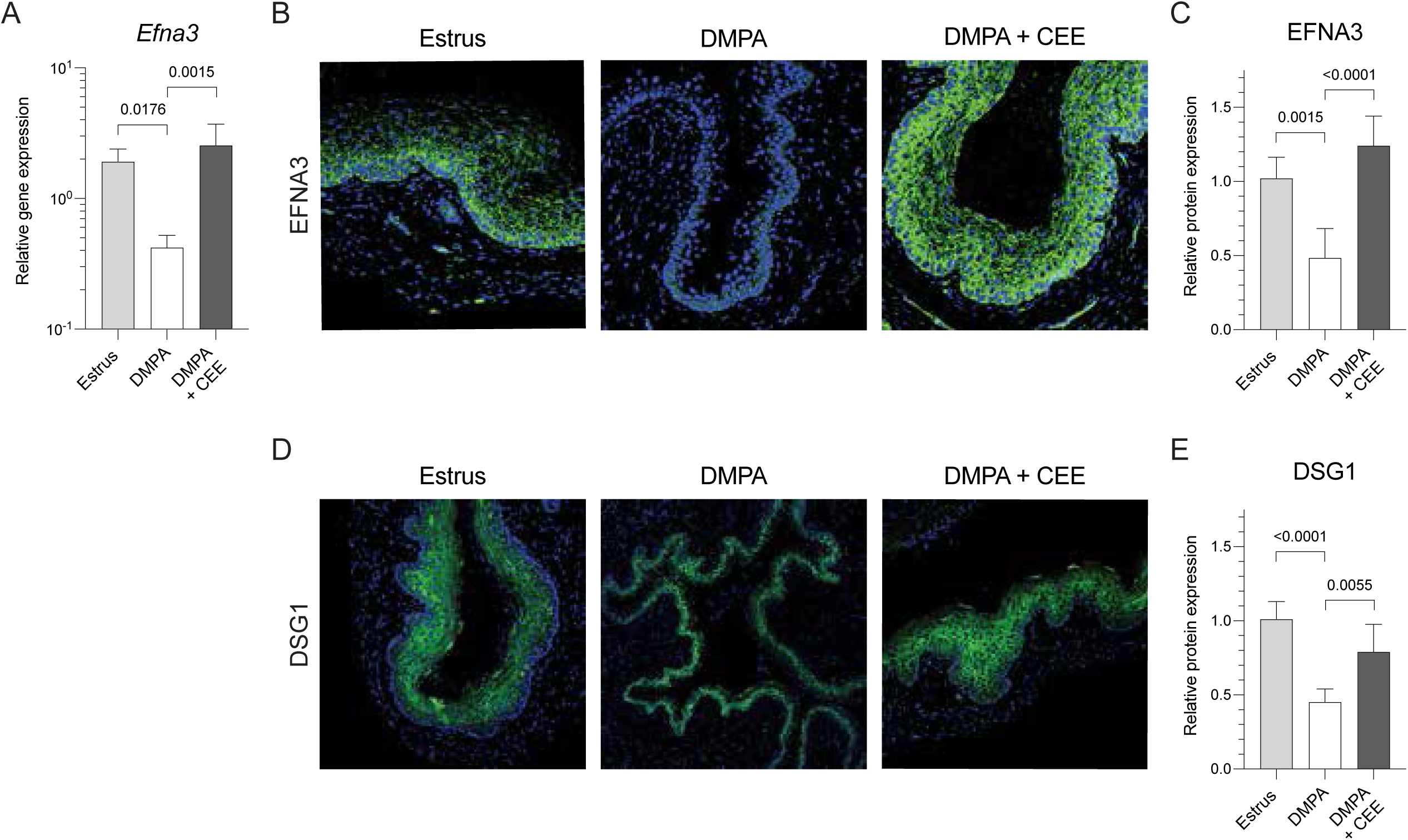
DMPA reduces vaginal expression of EFNA3 and DSG1 protein. Compared to estrus-stage mice or mice administered DMPA and CEE cream, DMPA-treated mice have significantly lower levels of EFNA3 and DSG1 protein. A) qRT-PCR quantifies lower ephrin-A3 (*Efna3*) gene expression in DMPA-treated vs. estrus-stage mice or mice treated with DMPA and CEE cream. B) Representative images from AF488-conjugated anti-mouse EFNA3 polyclonal antibody staining of vaginal tissue. C) Bar graph shows significantly lower levels of EFNA3 protein in DMPA treated vs. estrus-stage mice or mice treated with DMPA and CEE cream. D) Representative images of immunofluorescence staining for DSG1 protein in these same 3 groups of mice. E) Bar graph identifies reduced levels of DSG1 protein in DMPA-treated vs. estrus-stage mice or mice treated with CEE cream and DMPA. CEE; conjugated equine estrogen.

### rEFNA3 promotes vaginal epithelial integrity and barrier function

Upon identifying involvement of the EFNA3 signaling pathway in exogenous sex steroid-mediated regulation of desmosomal cadherin expression in mouse vaginal epithelium, we explored the ability of two forms of rEFNA3 (i.e., Fc fusion protein or His-tagged protein) to restore the loss of vaginal epithelial integrity in DMPA-treated mice. We elected to use mouse rEFNA3 Fc fusion protein (hereafter termed Fc tag) for its stability and mouse rEFNA3 His-tagged protein (hereafter termed His tag) as it more closely resembled the native cell surface protein. In initial studies, we intravaginally administered various doses of Fc tag or His tag (in PBS solution) to DMPA-treated mice. Other DMPA-treated mice received PBS with Fc control or PBS alone. Compared to Fc control- or PBS-treated controls, mice that received 1 μM of Fc tag or His tag displayed higher vaginal levels of DSG1 protein (Fig. 3A-3D). The 2 rEFNA3 treatments shared a biphasic dose-dependent effect, as concentrations greater than 1 μM were associated with lower levels of vaginal DSG1 protein (Fig. 3A-3D). Also compared to controls, DMPA-treated mice topically administered 1 μM of Fc tag or His tag exhibited greater vaginal epithelial thickness (Fig. 3E). Together, these studies identified intravaginal rEFNA3 administration reduces the loss of vaginal epithelial thickness and integrity generated by systemic DMPA injection.

**Figure 3.**
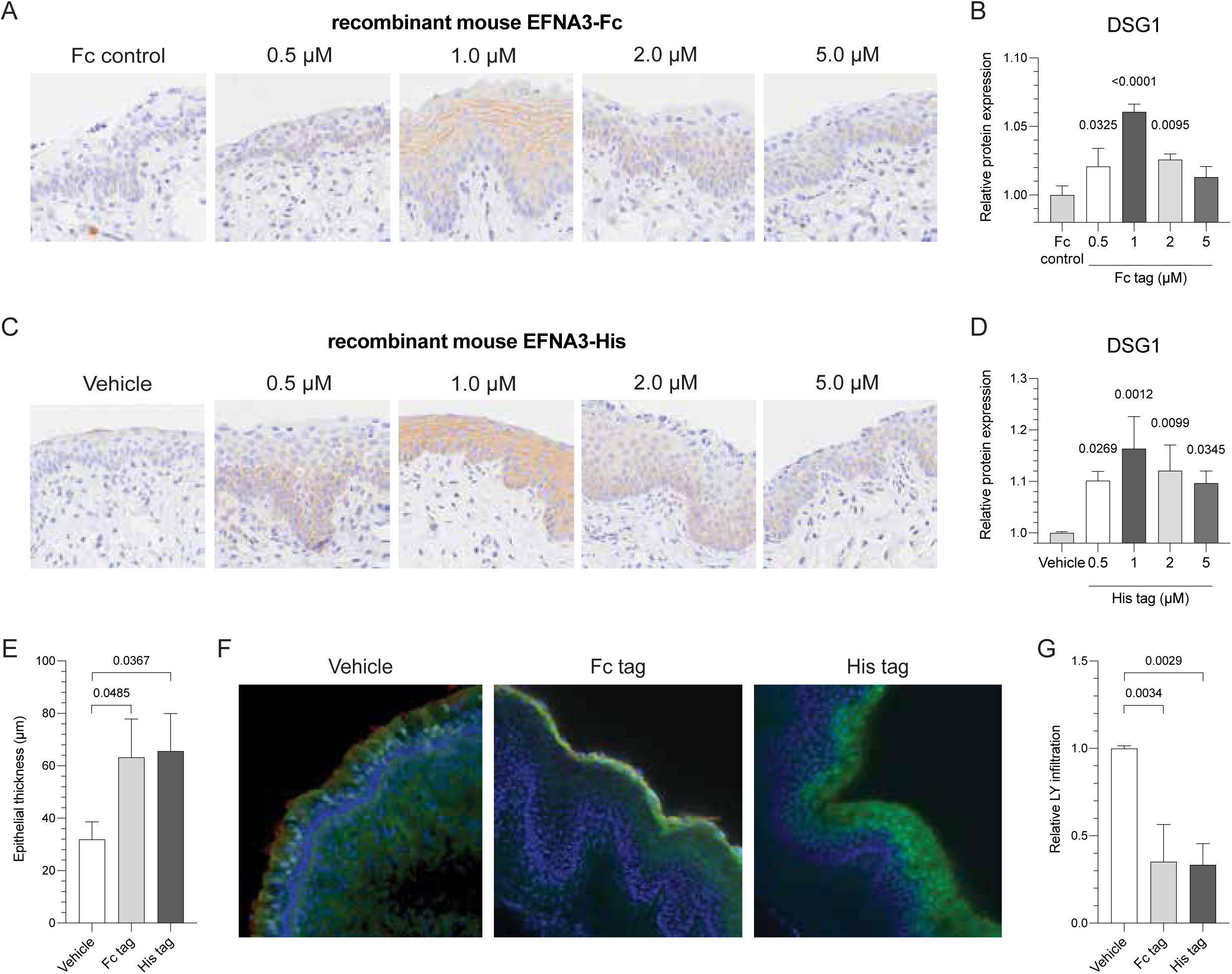
Topical recombinant EFNA3 (rEFNA3) increases DSG1 and DSC1. A) Representative images of DSG1 immunohistochemical staining 24 hours after DMPA-treated mice were intravaginally administered PBS with Fc control or indicated concentrations of Fc tag. B) Column graph identifies the biphasic dose-dependent effect on DSG1 protein levels upon treatment with Fc tag. C) Representative images of DSG1 immunohistochemical staining 24 hours after DMPA-treated mice were intravaginally administered PBS (Vehicle) or indicated concentrations of His tag. D) Column graph reveals comparable biphasic dose-dependent manner of Fc tag effect on DSG1 protein levels. E) Vaginal epithelial thickness 24 hours after DMPA-treated mice received PBS or 1 μM of Fc tag or His tag. F) Representative confocal microscopy images of vaginal epithelial permeability to a LMM fluorescent molecule (LY) intravaginally administered 24 hours after DMPA-treated mice received PBS or 1 μM of Fc tag or His tag. G) Bar graphs depict reduced LY penetration in vaginal epithelium of mice receiving 1 μM of either rEFNA3. Fc tag, mouse rEFNA3 Fc fusion protein; His tag, mouse His tag containing rEFNA3.

As rEFNA3 reduced the loss of vaginal integrity in DMPA-treated mice, we explored the ability of rEFNA3 to promote vaginal epithelial barrier function and restrict penetration of LY into the vaginal lamina propria of DMPA-treated mice. While LY readily penetrated vaginal epithelium to enter the lamina propria of DMPA-treated controls, it only penetrated superficial vaginal epithelial layers in mice administered DMPA and 1 μM of Fc tag or His tag (Fig. 3F, 3G). These results suggested that topical rEFNA3 lessened DMPA-mediated loss of vaginal epithelial barrier function. As ligand-receptor promiscuity is a recognized feature of Eph/EFN signaling pathways ^33^, we also explored ability of topical treatment with recombinant His-tagged versions of other EFNA to diminish DMPA-mediated losses of vaginal epithelial integrity and barrier function. While 1 μM of rEFNA2 promoted expression of the *Dsg1* and *Dsc1* genes and DSG1 protein more effectively than rEFNA1, rEFNA4 or rEFNA5, it was less effective than rEFNA3 at improving these parameters (Fig. 4A-4D). Also compared to rEFNA2, topical administration of rEFNA3 more effectively recovered vaginal epithelial barrier function and better restricted the penetration of LY into the vaginal epithelium (Fig. 4E, 4F). On the other hand, neither rEFNA2 nor rEFNA3 treatment approached the effectiveness of topical CEE in restoring the loss of vaginal epithelial barrier function generated by DMPA (Fig. 4C-4F).

**Figure 4.**
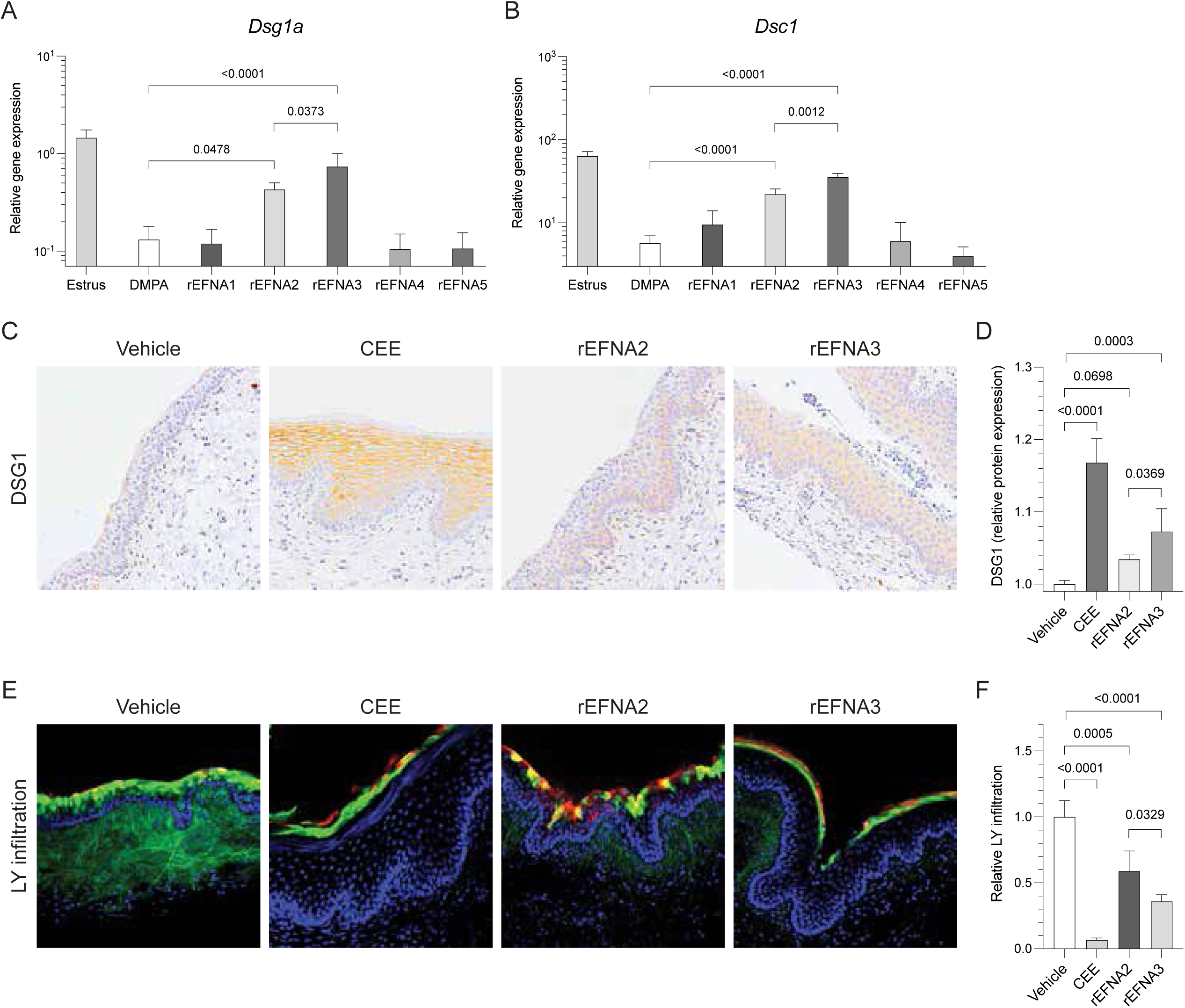
Vaginal epithelial barrier function is more effectively restored by rEFNA3 vs. rEFNA2. qRT-PCR quantifies *Dsg1a* (A) and *Dsc1* (B) expression after indicated rEFNA were administered to DMPA-treated mice. C) Representative images of DSG1 immunohistochemical staining induced by rEFNA2 vs. rEFNA3. D) Bar graphs display increased vaginal DSG1 levels in DMPA-treated mice receiving rEFNA2 or rEFNA3. E) Representative confocal microscopy images display reduced penetration of LY into vaginal epithelium of DMPA-treated mice that received rEFNA3 vs. rEFNA2. F) Bar graph identifies more effective recovery of vaginal epithelial barrier function in DMPA-treated mice administered rEFNA3 vs. rEFNA2.

### rEFNA3 delays HSV-2-induced morbidity

In DMPA-treated mice, genital HSV-2 infection is uniformly lethal as virus enters the central nervous system to cause fatal encephalopathy ^17, 34^. Because 1 μM of Fc tag or His tag treatment only partially restored vaginal epithelial barrier function in DMPA-treated mice, we hypothesized that treatment with either recombinant protein delays morbidity onset but does protect DMPA-treated mice from genital HSV-2 infection. As posited, HSV-2-mediated genital pathology appeared sooner in DMPA-treated mice vs. DMPA-treated mice topically administered 1 μM of rEFNA3 one day prior to infection. Genital pathology developed significantly faster in DMPA-treated mice, with no mice administered rEFNA3 exhibiting pathology prior to dpi 5 (Fig. 5A-B). Consistent with the appearance of genital pathology that was delayed, but not prevented, all rEFNA3-treated mice succumbed to HSV-2 (median survival was 9.5 days for either treatment group vs. 6 days for DMPA-treated mice). Conversely, all mice that received DMPA and CEE were protected from genital HSV-2 infection (Fig. 5C).

**Figure 5.**
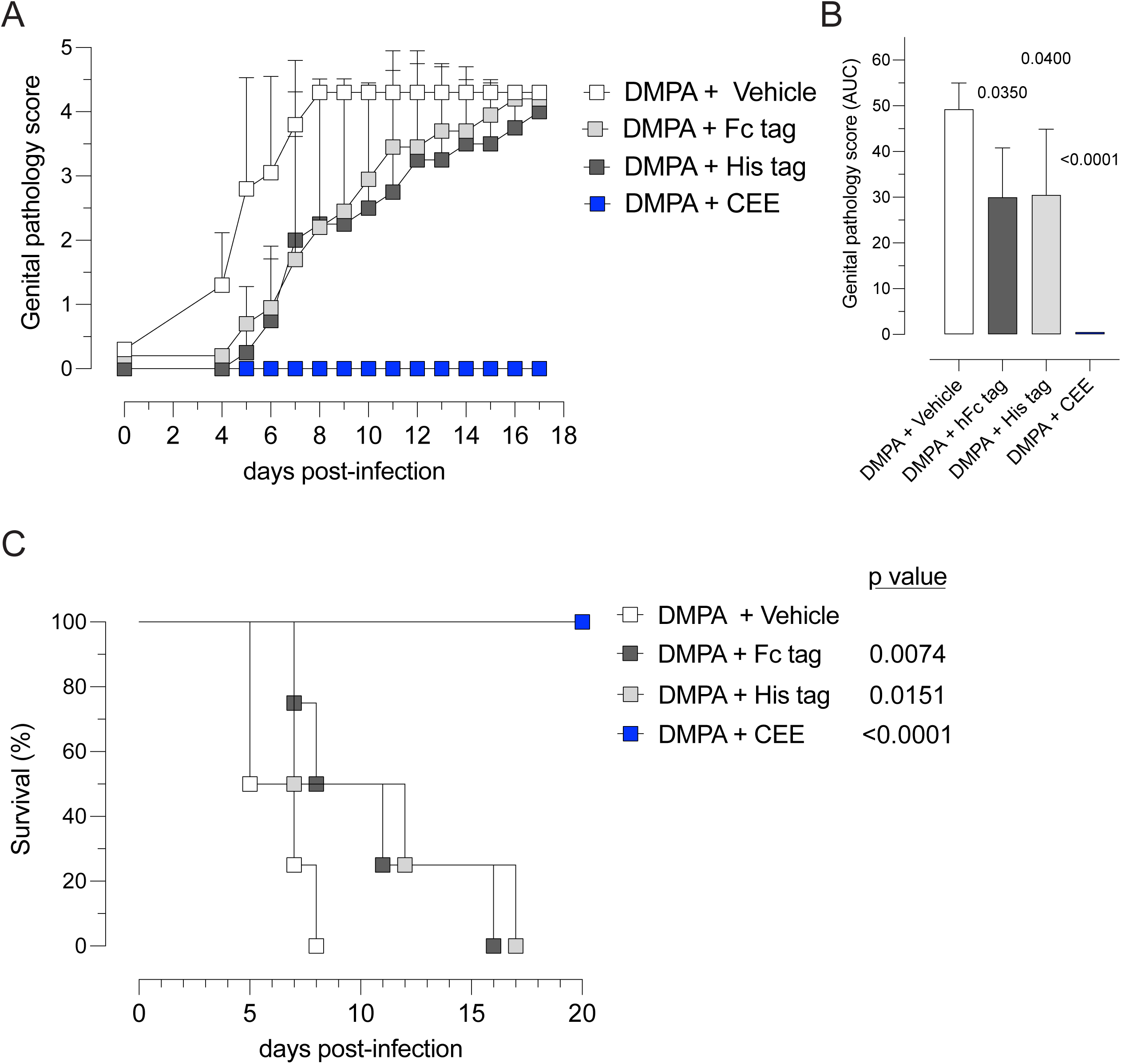
rEFNA3 delays onset of HSV-2-induced morbidity. DMPA-treated mice were administered a single 1 μM dose of Fc tag or His tag in PBS or PBS alone 24 hours prior to genital infection with 10^4^ pfu of HSV-2. Other DMPA-treated mice received topical CEE cream 24 hours before HSV-2 infection. All mice were evaluated daily post-infection. A) Cumulative genital pathology scores indicate accelerated onset of genital morbidity in controls vs. mice administered either form of rEFNA3. B) Column graph that provides comparison of the area under the curve for each mouse’s course of genital pathology. C) Survival curve identifies delayed onset of mortality in mice administered rEFNA3 vs. PBS alone. No mice that received CEE cream succumbed to infection.

## DISCUSSION

Earlier research established that desmosomes or anchoring junctions maintain barrier function in stratified squamous cutaneous epithelium ^35^ and that profound loss of barrier protection occurs in humans and mice genetically deficient in DSG1 ^6–8^. In addition, individuals genetically DSG1 deficient also display increased susceptibility to allergens and recurrent cutaneous infection ^6–8^. Indicative of their role in barrier protection, increased DSG1 and DSC1 expression is seen in more superficial layers of cutaneous epithelium ^36^.

Previously, we identified an analogous role for the desmosomal cadherins DSG1 and DSC1 in stratified squamous genital epithelium of humans, nonhuman primates, and mice ^11, 13–15, 18^. These reports newly identified that endogenous and exogenous sex steroids impact genital epithelial barrier function by altering genital DSG1 and DSC1 levels ^11, 14^. More specifically, we showed that higher levels of endogenous E in estrus-stage mice or topical treatment with CEE are associated with increased vaginal levels of DSG1 and DSC1 and robust vaginal epithelial barrier function whereas opposite effects in vaginal epithelium are generated in mice treated with the synthetic progestin DMPA ^11, 14^. The current investigation confirms and extends these results, performing genome-wide gene expression profile analysis and qRT-PCR studies that detected lower levels of *Dsc1* and *Dsg1*, IHC studies that measured reduced levels of DSG1 protein, and confocal microscopy studies that visualized the loss of barrier function in vaginal epithelial tissue from DMPA-treated mice vs. estrus-stage mice or mice administered DMPA and topical CEE. Consistent with prior clinical reports of increased genital inflammation in women using DMPA ^37, 38^, our gene expression analyses further identified that DMPA treatment activated inflammatory pathways associated with neutrophil degranulation and IL-10 signaling. Providing mechanistic explanation that couples DMPA-mediated inflammation with loss of barrier protection, we previously showed that DMPA treatment does not promote vaginal inflammation in germ-free mice (i.e., mice that lack vaginal microbiota) ^11, 39^. In other words, it seems likely that the DMPA-mediated loss of epithelial barrier protection promotes inflammatory responses by facilitating entry of endogenous host microbiota into vaginal tissue.

Our investigation also newly recognizes that DMPA suppresses several EFNA-driven pathways and, more specifically, that EFNA3 signaling regulates the expression of desmosomal cadherins that promote integrity and barrier protection in vaginal epithelial tissue. Erythropoietin-producing hepatocellular (Eph) receptors are a large family of receptor tyrosine kinases that are divided into A and B classes based on binding affinity and sequence homology ^40^. These receptors promiscuously bind ephrin ligands with variable affinity. For example, nine EphA receptors bind five ephrin-A (EFNA) ligands that have single receptor-binding domains bound to plasma membrane via GPI anchors ^41–44^. Consistent with our current results using murine vaginal epithelial tissue, prior studies identified that bidirectional EFNA-EphA signals regulate epithelialization and that EFNA3 is an upstream regulator of DSG1 and DSC1 expression in cutaneous epithelium ^28, 29^. Of note, cutaneous and vaginal epithelium are composed of basal (stratum basale), suprabasal (stratum spinosum and stratum granulosum), and cornified (stratum corneum) layers. While EphA receptors are expressed in each layer, EFNA ligands are concentrated in basal layers sites of more active epithelial cell proliferation ^29^. Conversely, soluble EFNA3 ligands may activate EphA receptors and trigger pathway signaling in EphA-expressing cells in more superficial layers of epithelium ^41–44^. Prior reports also identified that E-mediated activation of estrogen receptor 1 stimulates epithelial cell proliferation in both suprabasal and superficial layers of vaginal ^45–47^, a finding consistent with the ability of topical CEE treatment to restore vaginal DSG1 levels in DMPA-treated mice and create strong epithelial barrier protection.

In addition to identifying EFNA3 as an important regulator of vaginal epithelial desmosomes, the current study shows that topical rEFNA3 treatment partially restores the DMPA-mediated loss of vaginal epithelial function. This result is congruent with earlier reports that showed Eph/Ephrin signaling pathways are key upstream regulators of DSG1 and that EFNA-mediated EphA activation increases keratinocyte proliferation and differentiation by upregulating expression of DSG1 and DSC1 ^27–29^. Though we saw rEFNA3 partially restore vaginal epithelial DSG1 levels and barrier function in DMPA-treated mice, unlike CEE treatment, it failed to protect these mice from HSV-2 infection. One possible explanation for this failure to protect is that as few as one HSV-2 virion may be sufficient for productive infection and virus dissemination to the central nervous system ^17, 34^. Alternatively, the rEFNA3 we used may ineffectively penetrate into deeper epithelial layers. The rEFNA3 used in this study had a molecular mass around 35 kDa and are considerably smaller than the 70 kDa Texas Red-dextran molecules that display limited ability to enter intact vaginal epithelium (Fig. 3). Offering a third potential explanation for the inability to prevent HSV-2 transmission, rEFNA3 may cause EphA2 receptor overactivation that impairs its ability to restore vaginal epithelial barrier function in DMPA-treated mice. Consistent with this possibility, EphA receptors were shown to undergo endocytosis upon ligand binding and receptor activation promotes degradation that extinguishes signaling ^48^. Of note, EFNA3 binds EphA2 receptors with high affinity and *in vitro* stimulation of ligand for longer than 15 minutes promotes EphA2 degradation ^49^. Consistent with these *in vitro* findings, we saw a biphasic dose-dependent effect with *in vivo* treatment, with rEFNA3 doses greater than 1 μM correlating with lower levels of DSG1 in vaginal epithelial tissue (Fig. 3).

While not the primary objective of the current investigation, our results indicate that rEFNA3 may offer the foundation for a nonsteroidal and nonhormonal approach to promoting or restoring epithelial barrier function in the female genital tract. As rEFNA3 only partially recovered vaginal epithelial barrier function in DMPA-treated mice (Fig. 5), further studies are needed to define the ability of more frequent treatment or different delivery methods of rEFNA3 to elicit more robust responses or to develop EFNA3 analogs with greater or more durable bioactivity. Developing this approach would have the potential to restore vaginal epithelial health in reproductive age women using progestin-only contraceptives and postmenopausal women with vulvovaginal atrophy, as both groups exhibit loss of vaginal epithelial integrity and barrier function ^11, 16^. As a potential early-step in this process, the current investigation identifies EFNA3 as an important regulator of desmosomal function in vaginal epithelium and delivers vital new understanding of the sex steroid-mediated mechanisms that regulate vaginal epithelial integrity and barrier function.

## Acknowledgments

The authors thank Nirk Quispe Calla, MD and the Ohio State University Biomedical Informatics Shared Resources for Bioinformatics Services for technical assistance.

## Author contributions

TLC and RDVM were responsible for study conception, experimental design, and data analysis. KA and ML performed experiments and data analysis. ML composed the initial manuscript draft. All authors contributed to discussions that shaped content of the submitted manuscript.

